# Quantifying individual differences in brain morphometry underlying symptom severity in Autism Spectrum Disorders

**DOI:** 10.1101/430355

**Authors:** Emmanuel Peng Kiat Pua, Gareth Ball, Chris Adamson, Stephen Bowden, Marc L Seal

## Abstract

The neurobiology of heterogeneous neurodevelopmental disorders such as autism spectrum disorders (ASD) are still unclear. Despite extensive efforts, most findings are difficult to reproduce due to high levels of individual variance in phenotypic expression. To quantify individual differences in brain morphometry in ASD, we implemented a novel subject-level, distance-based method on subject-specific attributes. In a large multi-cohort sample, each subject with ASD (n=100; n=84 males; mean age: 11.43 years; mean IQ: 110.58) was strictly matched to a control participant (n=100; n=84 males; mean age: 11.43 years; mean IQ: 110.70). Intrapair Euclidean distance of MRI brain morphometry and symptom severity measures were entered into a regularised machine learning pipeline for feature selection, with rigorous out-of-sample validation and bootstrapped permutation testing. Subject-specific structural morphometry features significantly predicted individual variation in ASD symptom severity (19 cortical thickness features, p=0.01, n=5000 permutations; 10 surface area features, p=0.006, n=5000 permutations). Findings remained robust across subjects and were replicated in validation samples. Identified cortical regions implicate key hubs of the salience and default mode networks as neuroanatomical features of social impairment in ASD. Present results highlight the importance of subject-level markers in ASD, and offer an important step forward in understanding the neurobiology of heterogeneous disorders.

## Introduction

The autism spectrum disorders (ASD) are a group of neurodevelopmental conditions characterized by impairments in social communication and restricted and repetitive behaviours (Wing, 1997). Definitive neurobiological mechanisms underlying ASD or other heterogeneous neurodevelopmental disorders have yet to be clearly delineated due to significant heterogeneity within and between individuals (Hahamy, Behrmann, & Malach, 2015). Magnetic Resonance Imaging (MRI) offers an in vivo method to assay neurobiological abnormalities, and has led to some promising findings of brain dysfunction in ASD neuroimaging (Uddin, Dajani, Voorhies, Bednarz, & Kana, 2017). However, group differences in brain structure or function in ASD remain frequently misidentified because of high levels of variability between and within individuals, giving rise to poor reliability and reproducibility of findings (Ecker, 2017).

In addition to the large phenotypic variation in individuals with ASD, neuroimaging studies are confounded by a number of methodological factors related to differences in image acquisition sites, anatomical sex, IQ, as well as age-dependent perturbations of neurodevelopment (Pua, Bowden, & Seal, 2017). For example, age-related whole brain volume alterations (Lange et al., 2015), cortical thinning (Wallace, Dankner, Kenworthy, Giedd, & Martin, 2010), and atypical surface area development (Mensen et al., 2017) in ASD are associated with continuous shifts throughout the lifespan. Identifying altered neurodevelopmental trajectories in ASD becomes even more complex as cortical volume can be further delineated into separable sub-components of cortical thickness and cortical surface, each with distinct genetic influences on development (Panizzon et al., 2009). Previous reports of cortical measures of brain structural morphometry in ASD have been inconsistent, such as increased regional cortical thickness (Ecker et al., 2013; Hardan, Muddasani, Vemulapalli, Keshavan, & Minshew, 2006; Raznahan et al., 2013), decreased (Hardan, Libove, Keshavan, Melhem, & Minshew, 2009), or with significant cortical thinning in frontal, temporal or parietal regions (Hadjikhani, Joseph, Snyder, & Tager-Flusberg, 2005). Similarly, surface area in ASD has been reported to be increased (Hazlett et al., 2011), decreased (Ecker et al., 2013), or not significantly different from neurotypical peers (Raznahan et al., 2013; Wallace et al., 2013). Alterations in grey matter volume in ASD have also been reported to be driven by the absence of typical age-related cortical thinning (Smith et al., 2016). These mixed findings suggest that the expression of ASD in atypical brain structure is likely to differ between individuals with the condition, and across different age cohorts. Consequently, research efforts to identify consistent differences in the brain in individuals with ASD have remained inconclusive. Given the diverse nature of the condition, there is an increasing need for predictive brain-based markers that are sensitive to heterogeneity in the neurobiology and symptom expression of ASD (Jack & Pelphrey, 2017).

Emerging work suggests that individual-specific variation in brain architecture may be a critical factor underlying idiosyncrasies in ASD symptom characteristics (Chen, Nomi, Uddin, Duan, & Chen, 2017; Dickie et al., 2017). As conventional neuroimaging investigations typically rely on broad between-group comparisons without sufficient consideration of subject-specific effects, such approaches have been of limited yield in ASD research. Additionally, high-dimensional neuroimaging data with a large number of, often co-linear, features relative to small sample sizes pose further issues with increased risk of false positives. In particular for such a heterogeneous population as ASD, these challenges suggest that investigations of brain structure and function in ASD should incorporate appropriate subject-level modelling, with adequate consideration for common problems associated with large-scale high-dimensional neuroimaging data analysis (Bzdok & Yeo, 2017; Mwangi, Tian, & Soares, 2014).

Drawing from multi-disciplinary methodologies in ecology and twin modelling (Carlin, Gurrin, Sterne, Morley, & Dwyer, 2005), we developed a novel subject-level, distance-based method to test the hypothesis that neuroanatomical differences between subjects can explain individual differences in symptom severity. Using this approach on carefully matched case-control pairs at the individual rather than group level, we compared subject-specific differences in brain structural morphometry on MRI to associated intrapair differences in individual symptom severity. Specifically, we hypothesized that intrapair differences in cortical thickness and surface area features could predict individual variation in ASD symptom severity. By investigating relative individual differences within conservatively matched subjects, confounding effects related to inter-subject or cohort differences such as age, sex, intelligence and image acquisition site are also implicitly controlled for. Importantly, our approach implements well-validated and sophisticated machine learning and feature reduction techniques to ensure reproducibility of findings with reduced likelihood of false positives.

## Results

Using machine learning to predict subject-specific differences in symptom severity from differences in MRI features (Figure 1 and 2), we applied regularised linear regression with elastic net penalty to achieve a sparse solution and select important features from the full imaging dataset. After training, the model significantly predicted differences in symptom severity between cases and controls in the out-of-sample dataset (*R*^2^=0.153; *p*=0.01, 5000 permutations). Based on 1000 iterations of the training loop, nineteen cortical thickness features were retained as predictors of individual differences in symptom severity (Figure 3A; Supplementary Table 1). The above procedure was repeated for cortical surface area measurements to predict differences in symptom severity. Ten surface area features were selected in the training dataset, with a favourable out-of-sample model fit (*R*^2^=0.18, *p*=0.006; Figure 3B; Supplementary Table 2).

**Table 1.** Descriptive statistics of matched samples

| **Group** | **n** | **Sex** | **Age** | **IQ** | **SRS** |
| --- | --- | --- | --- | --- | --- |
| ASD | 100 | n=84<br>males | 11.45 (3.51);<br>Range: 5.92-24.58 | 110.58 (13.18);<br>Range: 80-149 | 91.7 (28.42);<br>Range: 11-162 |
| Controls | 100 | n=84<br>males | 11.43 (3.55);<br>Range: 5.89-23.92 | 110.70 (13.18);<br>Range: 79-148 | 19.7 (13.0);<br>Range: 1-57 |

*Note. SRS: Social Responsiveness Scale raw scores. Higher score indicates more severe ASD symptoms.*

**Figure 1.**
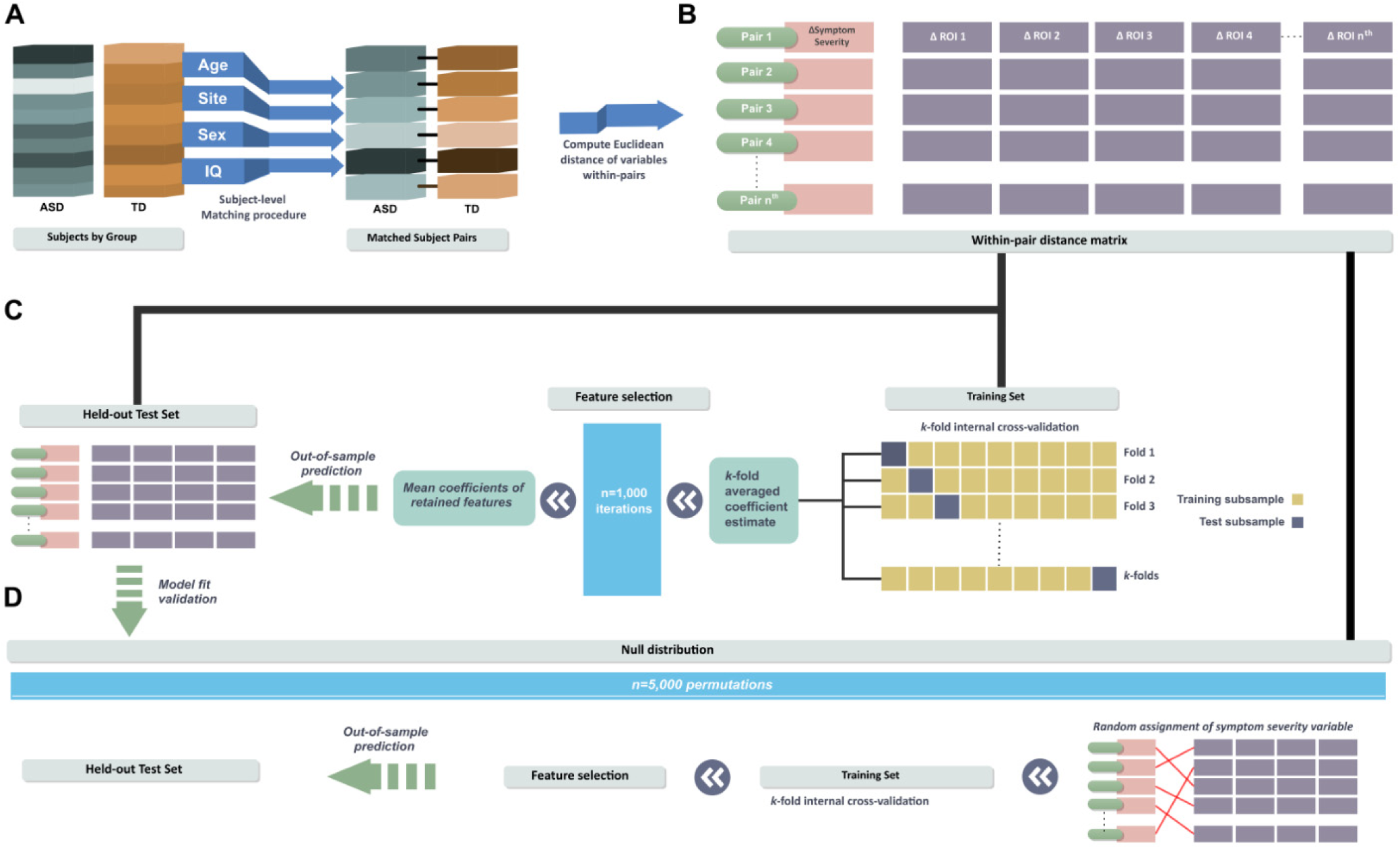
Subject-level distance-based pipeline. **A:** Each ASD case was individually matched to one control participant in age, sex, IQ and acquisition site. **B:** For every matched pair, within-pair Euclidean distances (δ) on symptom severity variables and morphometry of brain region-of-interests (ROI) were computed. **C:** Using a machine learning approach, regularised regression with elastic net penalization was implemented to test the association between within-pair δROI and δsymptom severity. A subset of the sample (33%) was held out as an independent out-of-sample test set. Remaining data was used as a training set to obtain cross-validated model weights for feature selection. The trained coefficient weights were then used to generate predictions and model fit parameters in the held-out test set. **D:** Finally, out-of-sample model

**Figure 2.**
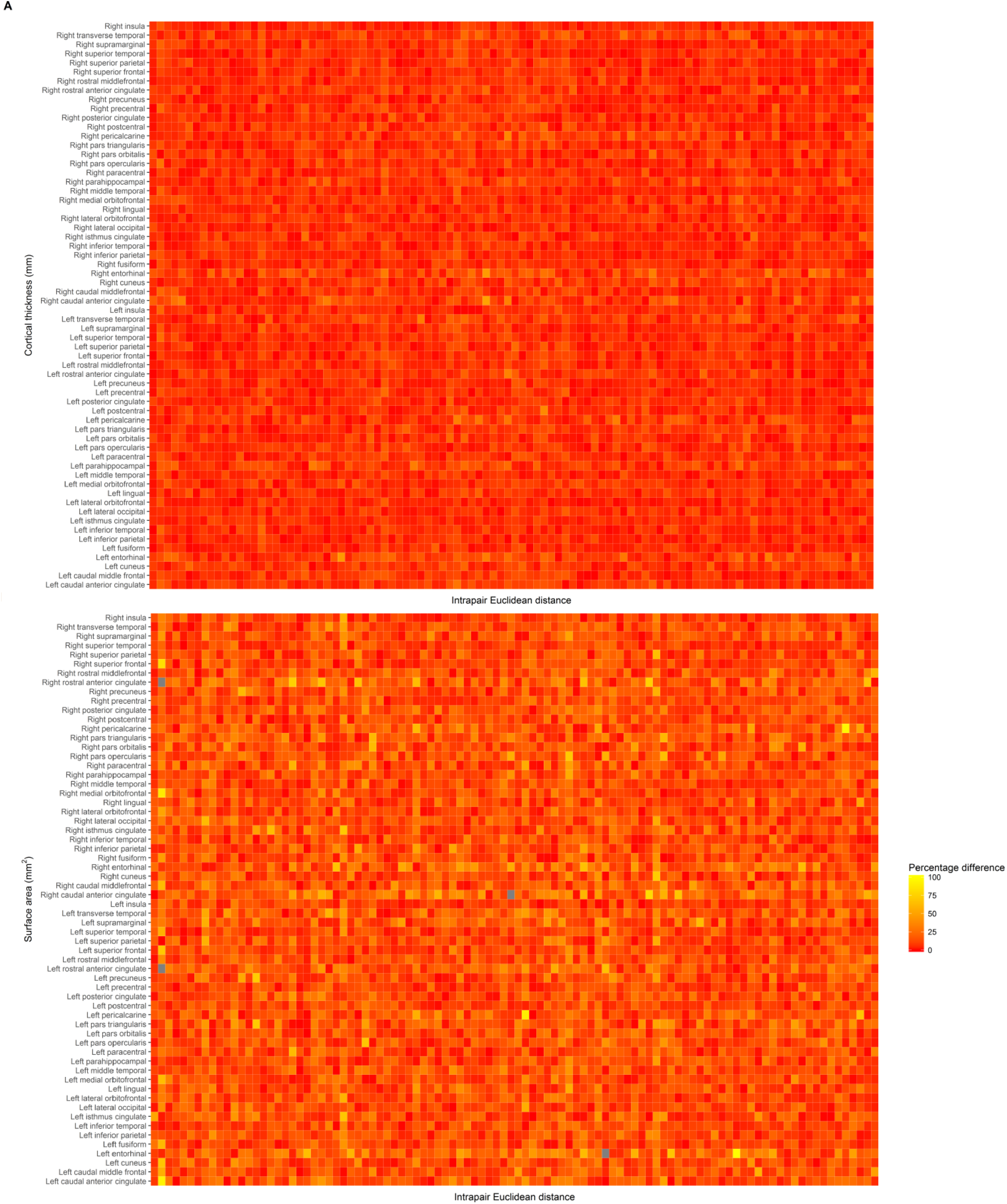
Exploratory data visualisation of intrapair differences for cortical thickness (A) and surface area (B). Rows represents the within-pair percentage difference in each cortical region (columns). Brighter heatmap colours (yellow) indicates higher intrapair difference in structural morphometry features. Neutral regions (grey) indicate a percentage difference exceeding 100%. Neutral regions (grey) indicate a percentage difference exceeding 100%. Subject-level differences in structural morphometry of specific regions, such as the anterior cingulate, appear to be higher than other cortical regions across most subjects.

**Figure 3.**
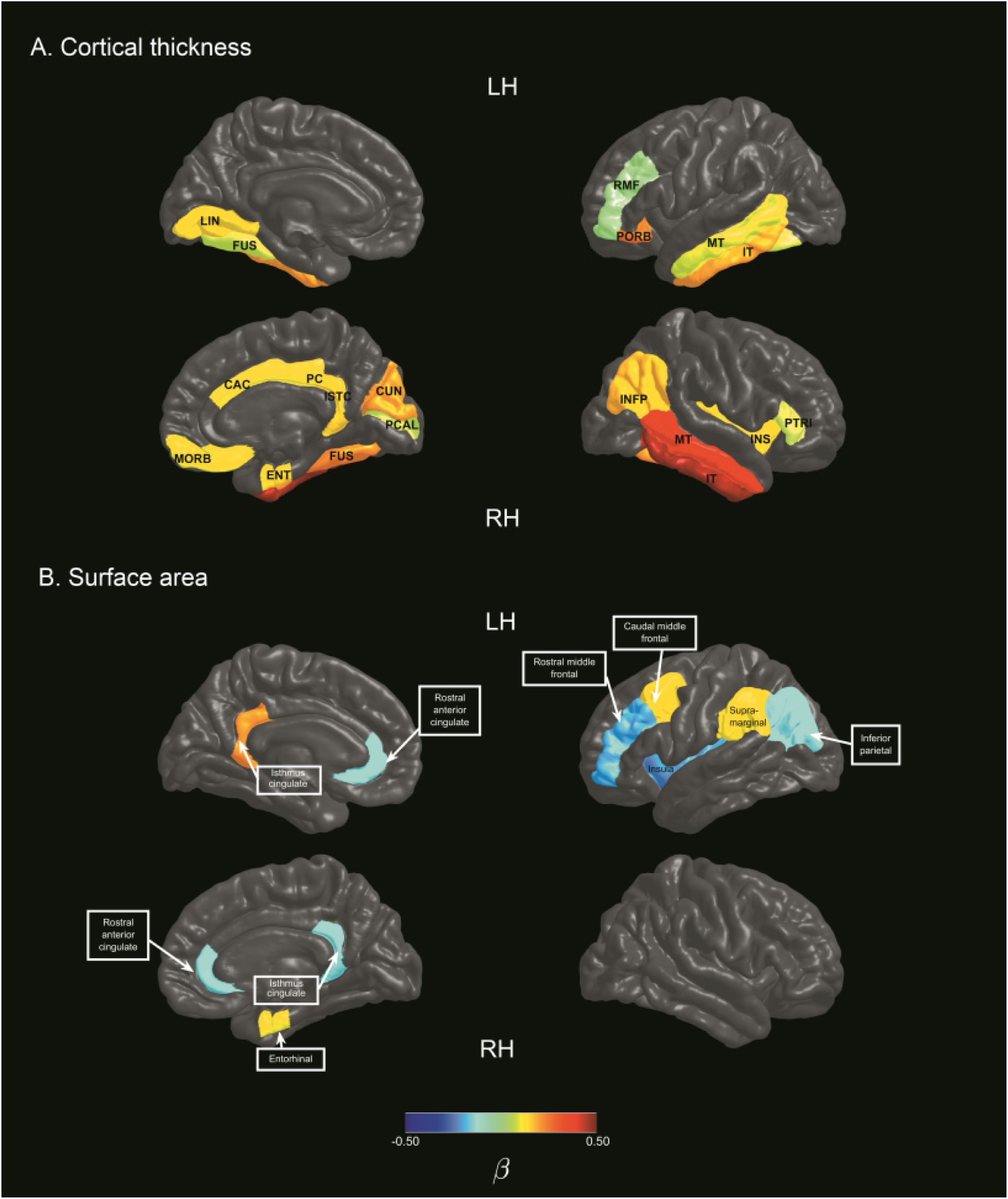
Cortical features selected using regularised regression models. Colour bars represent mean beta coefficients of cortical regions associated with individual differences in symptom severity in ASD. **A:** Cortical thickness features associated with symptom severity variation in ASD **B:** Surface area features associated with symptom severity variation in ASD. ***Note.*** *CAC: caudal anterior cingulate gyrus; CUN: cuneus; ENT: entorhinal; FUS: fusiform gyrus; INFP: inferior parietal gyrus; INS: insula; ISTC: isthmus cingulate gyrus; IT: inferior temporal gyrus; LH: left hemisphere; LIN: lingual gyrus; MORB: medial orbitofrontal; MT: middle temporal gyrus; PCAL: pericalcerine; PC: posterior cingulate gyrus; PORS: pars orbitalis; RH: Right hemisphere; PTRI: pars triangularis; RMF: rostral middle frontal gyrus.*

We validated our approach against more conventional methods of group-level prediction less robust to heterogeneity across individuals. As expected, without accounting for subject-level within-pair differences, regression model training to predict symptom severity based on group-level MRI features demonstrated a poor fit in out-of-sample validation tests (Cortical thickness model fit: *R*^2^= 0.0000434; surface area model fit: *R*^2^=0.0158) by comparison. Comparisons of model fit performance for both approaches are shown in Supplementary Figure 1 and Supplementary Table 3.

## Discussion

By implementing a strict matching procedure combined with subject-level distance-based prediction of variation in ASD symptom severity, we demonstrated that individual-specific differences in cortical morphology were associated with subject-level variation in ASD symptom severity. Key cortical features implicate abnormal morphometry of frontal and temporal-parietal cortices, fusiform gyri, anterior and posterior cingulate regions, and the insula.

Cortical surface area features identified in the present study were strikingly consistent with previous findings of altered surface area underlying increased grey matter volume in 3-year old boys with ASD (Ohta et al., 2016). Cortical surface area in 8 of the 10 cortical regions identified in the present study (Supplementary Table 2) were reported by Ohta and colleagues to be significantly increased in ASD compared to controls, with the exception of the bilateral isthmus cingulate. Specific regions reported in both studies were the left caudal middle frontal, left rostral middle frontal, right entorhinal, left inferior parietal, left supramarginal, bilateral rostral anterior cingulate, and the left insula. Given the known inconsistencies across investigations on ASD structural morphometry, the similarity in reports of atypical surface area features across both studies that independently identified the same set of cortical regions and laterality is remarkable. Compared to the study of Ohta et al. (2016) that investigated males at the age of 3 years, present findings were derived from independent samples of an older age cohort (age range 5 – 25 years) that also included females, and utilised a different analysis method with the novel subject-level distance-based approach. The consistent results across different age cohorts and samples suggest that differences in the identified surface area features may be a stable feature of ASD over time.

Similarly, Wee, Wang, Shi, Yap, and Shen (2014) reported that morphological abnormalities in a set of cortical regions were the most discriminative features for the classification of ASD between 5 to 23 years of age. Using a multi-kernel learning strategy for feature selection, classification based on regional and interregional features in unseen samples achieved a sensitivity of 95.5% and specificity of 97%, with an accuracy of 96.27% and area under receiver operating characteristic curve (AUC) of 0.995, suggesting acceptable predictive utility. Similar cortical regions underlying individual differences in ASD were identified in the present study in the surface area of the left caudal middle frontal, left supramarginal, right rostral anterior cingulate gyrus, and cortical thickness of the right inferior temporal gyrus, right cuneus, left middle temporal and right fusiform gyrus.

Other neuroimaging investigations in ASD have also implicated the identified cortical regions in either or both hemispheres in the anterior cingulate (Haznedar et al., 1997; Jiao et al., 2010; Prigge et al., 2018), posterior cingulate (Hyde, Samson, Evans, & Mottron, 2010; Prigge et al., 2018; Yang, Beam, Pelphrey, Abdullahi, & Jou, 2016), isthmus cingulate (Caeyenberghs et al., 2016; Doyle-Thomas et al., 2013; Yang et al., 2016), insula (Doyle-Thomas et al., 2013), rostral middle frontal gyrus (Hyde et al., 2010), pars orbitalis (Caeyenberghs et al., 2016), pars triangularis (Jiao et al., 2010), medial orbitofrontal (Hyde et al., 2010; Jiao et al., 2010), middle temporal gyrus (Abell et al., 1999; Yang et al., 2016), inferior temporal gyrus (Abell et al., 1999; Prigge et al., 2018; Yang et al., 2016), fusiform gyrus (Hyde et al., 2010), inferior parietal lobule (Hyde et al., 2010), supramarginal gyrus (Zielinski et al., 2014), lingual gyrus (Hyde et al., 2010; Prigge et al., 2018; Zielinski et al., 2014), cuneus cortex (Zielinski et al., 2014), and pericalcerine cortex (Prigge et al., 2018). Together with previous reports of similar clusters of cortical features (e.g. Ohta et al., 2016; Wee et al., 2014) identified in the present study, convergent results across multiple independent investigations suggest that atypical structural morphometry in these specific regions may be characteristic of altered neurodevelopment in ASD.

These distributed cortical regions facilitate key aspects of social, language, and sensory functioning, deficits of which are consistent with clinical features in ASD. For example, the middle temporal and inferior temporal gyri subserve language and semantic processing, and visual perception (Chao, Haxby, & Martin, 1999; Herath, Kinomura, & Roland, 2001). The right middle temporal gyrus and right insula are part of a distributed cortical network for modulating attention to salient features of the multimodal sensory environment (Downar, Crawley, Mikulis, & Davis, 2000). The right fusiform gyrus and occipital-temporal are highly specialised for face perception, recognition, and representation of facial features such as eye gaze and facial expressions that are necessary for social communication (Kanwisher, McDermott, & Chun, 1997; Rossion et al., 2003; Rossion, Schiltz, & Crommelinck, 2003). Notably, the identified cortical regions in the cingulate and insula implicate hub regions of the salience network (SN) and default mode network (DMN) that have been increasingly suspect to be aberrant in ASD (Anderson et al., 2011). The SN primarily anchored to the anterior insular and dorsal anterior cingulate cortex contributes to cognitive and affective processes such as social behaviour and communication, and the integration of sensory, emotional and cognitive information (Menon, 2015). The DMN comprising the posterior cingulate, medial prefrontal and parietal cortices typically demonstrates reduced activity on task initiation, and functions to support self-referential and introspective states and social cognition (Mak et al., 2017). Altered function of the salience network and DMN are highly consistent with ASD symptomatology, and emerging evidence suggests that specific features of cortical regions comprising these networks may discriminate ASD from neurotypical development (Uddin et al., 2017).

### A subject-level distance-based framework

Morphometry of the frontal, temporal, fusiform and insular cortices have been suggested to be important classification features in ASD. However, findings were inconsistent due to high variability in symptom severity, age and IQ across heterogeneous ASD subgroups. Classification accuracy became poorer as subgroup differences along these variables increased, but was significantly improved by matching subgroups on subject demographics (Katuwal, Baum, Cahill, & Michael, 2016). Between-group differences in cortical thickness in ASD have also been reported to become non-significant after controlling for IQ, further highlighting the importance of matching on key confound variables (Hardan et al., 2009). Present findings support the hypothesis of Katuwal and colleagues that increasing homegeneity between case and control populations can reduce noise and improve precision in classification. Further, similar cortical regions were also implicated in ASD in cortical thickness of occipital-temporal regions and the anterior cingulate. By modelling individual differences in well-matched subgroups, consistent patterns of abnormalities across cortical regions and subjects may emerge or become more distinct. For example, intrapair differences between cases and controls in structural morphometry of certain regions, such as the anterior cingulate, appear to exceed differences observed in other cortical regions in a large proportion of the sample (Figure 2). To qualify as potential candidate markers in the neurobiology of ASD however, such putative features must necessarily demonstrate significant associations with ASD symptomatology based on rigorous analysis and validation (Pua, Malpas, Bowden, & Seal, 2018). Using the subject-level distance-based framework in the current study, we demonstrated that individual differences in the structural morphometry of identified cortical regions also predicted subject-level variation in symptom severity (Figure 3).

Present results suggest that as neuroanatomy diverges between ASD and control subjects, individual differences in symptom severity increase within matched case-control pairs. Conversely, negative coefficient weights reflect an increase in symptom severity differences as regional cortical features becomes more similar between ASD and controls. While this may appear counterintuitive, the direction of differences in altered brain morphometry in ASD have been observed to shift across development, due to age-dependent changes in the neurodevelopmental trajectory of ASD that differs from controls (Lin, Ni, Lai, Tseng, & Gau, 2015). Previous longitudinal findings suggest that the expected typical age-related decline in cortical surface area (Mensen et al., 2017) or cortical thickness (Smith et al., 2016) may be absent in ASD. Group differences in cortical thickness also varied across development stages, with region-specific differences in age-related trajectories between ASD and controls (Zielinski et al., 2014). A dynamic pattern of age-specific abnormalities has also been reported with increased cortical thickness in children with ASD. However, no differences were observed in adult cohorts due to an accelerated rate of cortical thinning in ASD compared to controls (Khundrakpam, Lewis, Kostopoulos, Carbonell, & Evans, 2017). As a consequence of the different developmental trajectories between groups, atypical cortical developmental trajectories in ASD may intersect with that of typically developing peers at certain stages of development. At such time-points, group differences in abnormally developing cortical regions in ASD may not be detected cross-sectionally, despite a concomitant difference in symptom severity. The presence of group-dependent developmental trajectories that are dynamic over time could explain the high prevalence of inconsistent findings in ASD. With our proposed framework, we were able to identify distinct patterns of abnormalities associated with symptom severity in ASD that were not dependent on detecting mean differences at the group level. Notably, our approach based on cross-sectional data identified structural differences in ASD in the same regions reported in the longitudinal study of (Mensen et al., 2017) in surface area of the anterior cingulate, insula, supramarginal gyrus and inferior parietal lobe, and cortical thickness of the bilateral inferior temporal gyri.

Importantly, a dimensional approach based on continuous measures allows for a more precise quantification of sub-threshold ASD traits in individuals who do not meet the criteria for clinical diagnosis, otherwise known as the broader autism phenotype (Dawson et al., 2002). For example, a broader phenotype individual in the control group may display a high degree of similarity in symptom severity to an individual with a milder presentation of ASD (Bishop, Maybery, Wong, Maley, & Hallmayer, 2006). This is consistent with overlapping distributions of SRS scores in children with or without ASD observed in a large nation-wide population sample, such that a proportion of controls displayed higher SRS scores than individuals with ASD, and vice versa (Kamio et al., 2013). In the present study, a similar pattern is observed with overlapping symptom severity scores ranges between cases and controls. Investigating group differences in ASD based on group-averaged variables between cases and controls are thus likely to obscure important subject-specific effects, and could explain inconsistent findings from previous studies.

Indeed, present results suggest that subject-level modelling significantly outperforms group-difference methods of symptom severity prediction in ASD. We show that individual variability in brain morphometry and symptom severity can be modelled with a subject-level distance-based approach. Dimensional approaches to symptom measurement as such could be more effective in delineating the association between properties of the brain and symptom severity. Individual differences in neurodevelopment were also accounted for based on confound matching at the individual level (rather than group) to improve inter-subject homogeneity. With the robustness of present findings, such methodological considerations may be important for improved characterisation of heterogeneity in ASD brain morphometry that better reflects the continuous spectrum of symptom severity in this population.

### Future directions

In the current study, individual differences in cortical thickness between ASD and controls were more prevalent in the right hemisphere, with a left hemisphere bias for differences in surface area features. Cortical features that were implicated bilaterally were the surface area of the rostral anterior cingulate and isthmus cingulate, and cortical thickness of the middle temporal, inferior temporal and fusiform gyrus. ASD has been related to a loss or inversion of typical patterns of brain asymmetry or lateralisation, with abnormal asymmetry in brain morphometry and connectivity associated with symptom deficits in ASD (Conti et al., 2016; Floris et al., 2016; Herbert et al., 2002). For example, the study of Wee et al. (2014) noted significantly more abnormalities in the right hemisphere than the left in ASD, in agreement with previous reports of a right-hemisphere bias in brain structural and functional asymmetry in ASD (Chiron et al., 1995; Herbert et al., 2004; Wei, Zhong, Nie, & Gong, 2018). In contrast, a leftward lateralisation of abnormalities in ASD was reported for cortical thickness (Khundrakpam et al., 2017) and surface area (Dougherty, Evans, Katuwal, & Michael, 2016). Distinct patterns of lateralisation between different morphological features in ASD may be related to the independent growth trajectories of cortical thickness and cortical surface area, each regulated by discrete genetic mechanisms (Panizzon et al., 2009). Further, the direction of atypical asymmetry in structural morphometry has been shown to shift throughout development in ASD with decreasing leftward asymmetry with age, or differ between high and low functioning individuals with ASD (Dougherty et al., 2016; Khundrakpam et al., 2017). Mixed findings of structural asymmetry or direction of effects in ASD could thus be due to the diverse aetiology or distinct subtypes in the condition, as well as relative differences to control that vary across developmental stages due to altered developmental trajectories (Moreno-De-Luca et al., 2013). While we have shown that individual differences in specific cortical features are strongly implicated in ASD, the complex expression of age-dependent changes in ASD that are dynamic over time requires further investigation beyond cross-sectional studies.

Together, present results derived from rigorous testing and validation techniques suggest that subject-level variation in brain properties are important characteristics in the expression of ASD. Present findings were limited to cortical thickness and surface area features, and other properties across different measures of brain structure and function could account for larger proportions of variance in symptom severity. The robustness of the subject-level distance-based approach is nevertheless promising. Given that complex neurodevelopment conditions such as ASD likely stem from perturbations of anatomically distributed and interconnected neural systems, future applications of the subject-level distance-based approach to investigations of intrinsic brain networks may reveal more sophisticated insights into atypical neural mechanisms in ASD (Fornito, Bullmore, & Zalesky, 2017). As brain structural connectivity constrains the development of functional networks across the lifespan, validation across different imaging modalities will be necessary to elucidate distinct neurobiological mechanisms, such as network analysis of white matter microstructure and functional connectivity investigations (Grayson & Fair, 2017).

We strongly encourage continued multimodal subject-level distance-based investigations to further challenge this hypothesis in large multi-site cohorts. As we have shown, investigations in ASD must necessarily demonstrate generalizability beyond in-sample modelling, given the high levels of inter-and intra-individual heterogeneity in this population. It is likely that the reliable identification of neural correlates in ASD strongly depends on quantifying individual variation in the phenotypic expression of ASD or the broader autism phenotypes. Such individualised approaches will be important for the development of clinical applications to aid management or personalized intervention strategies for each unique patient at the individual level.

## Conclusion

The robustness and generalisability of present findings is important progress in the search for neural correlates of heterogeneous disorders such as ASD, and offers a promising insight into the neurobiology of ASD symptomatology. Based on present results with out-of-sample predictions, cortical hubs of the salience and DMN networks are likely to be implicated as potential neuroanatomical markers of ASD symptomatology. Increased reliability and validity of evidence for subject-specific alterations in brain structure and function will be necessary to advance current knowledge about the aetiology of ASD, where individual variability should be carefully modelled, rather than discarded as noise.

## Methods

### Participants

Data was obtained from the Autism Brain Imaging Database Exchange (ABIDE-II) cohort across 17 independent imaging sites (Di Martino et al., 2017). Protocols specific to each imaging site for diagnosis, behavioural and cognitive assessment, and Magnetic Resonance Imaging (MRI) acquisition are publicly available^1^. The Social Responsiveness Scale (SRS; Constantino & Gruber, 2012) was used as a phenotype measure of ASD symptom severity. The SRS has been established to be a reliable and valid quantitative measure of ASD traits, demonstrating convergent validity with the gold standard Autism Diagnostic Observation Schedule and Autism Diagnostic Interview, and is able to discriminate ASD from other psychopathologies (Bölte, Poustka, & Constantino, 2008; McConachie et al., 2015). The instrument is commonly used for both screening and as a tool to aid clinical diagnosis.

Based on the multivariate genetic matching method of Ho, Imai, King, and Stuart (2011), each ASD case was individually matched to the nearest control participant in age and IQ, restricted to a maximum distance of 0.25 standard deviations for each variable within each pair. For categorical variables, exact matching criteria were set for participant sex and image acquisition site. The genetic search algorithm (Diamond & Sekhon, 2013) aims to achieve optimal balance after matching by finding a set of weights for each covariate of interest. Matching balance was evaluated by univariate and paired t-tests for dichotomous variables and the Kolmogrov-Smirnov test for continuous or multinomial variables. The process selected n=100 individuals with ASD to 100 controls eligible for inclusion (see Table 1).

### Image processing and quality control

Pre-processing and analysis of T1-weighted (MRI) was performed using FreeSurfer (v6.0.0; http://surfer.nmr.mgh.harvard.edu/). Visual inspection for movement artefacts were conducted for every subject to ensure data quality control. We inspected images for characteristic ring artefacts caused by in-scanner head motion, and evaluating grey/white matter and grey matter/CSF boundaries. Briefly, the FreeSurfer cortical surface reconstruction pipeline performs the following steps in sequence: non-uniform intensity correction, skull stripping, segmentation into tissue type and subcortical grey matter structures, cortical surface extraction and parcellation. Manual editing was performed on the white matter mask images to avoid false-positive errors in estimated surfaces and to ensure accurate masking of the dura. We further inspected images for topological errors during cortical reconstruction by inspecting surface outputs and parcellations for each subject. Cortical morphometric statistics based on the Desikan–Killiany–Tourville (DKT) brain anatomical atlas were used to estimate cortical thickness (mm) and surface area measurements (mm^2^) for each brain region. An advantage of the DKT atlas is that labelling is performed on a per-subject basis rather than the projection of a single parcellation onto every cortex, and is well-suited for the present subject-level analyses. Given that participants with high motion in one scan are also likely to move more in other scans during the same session (Savalia et al., 2017), we further evaluated head motion parameters in each subject’s resting-state functional MRI (fMRI) scan using framewise displacement (FD) as an estimate of volume-to-volume head movement. Inter-subject mean FD was not significantly associated with differences in SRS scores within matched subject pairs (t=1.54, p=0.125), suggesting that individual variation in symptom measures were not likely explained by differences in in-scanner head motion.

### Subject-level distance-based analysis

Within-pair Euclidean distances were computed for every matched pair on clinical (SRS score) and demographic (age and IQ) continuous variables and brain structural morphometry measures (cortical thickness and surface area). Age and total intracranial volume were regressed out using standardised residual adjustment to adjust for inter-subject disparity in age and head size (O’Brien et al., 2011). By measuring the anatomical difference between matched brains at the level of individual subjects, we can investigate brain-behaviour associations underlying symptom severity by testing whether differences in brain structure can explain differences in symptom severity between paired subjects. Subjects were also strictly matched at the individual level on key neurodevelopmental and demographic factors such as age, sex, IQ, and MRI acquisition site. This improves subgroup equivalence (Stout, Wirtz, Carbonari, & Del Boca, 1994), and reduces the likelihood of significant differences on these confound covariates influencing observed outcomes between subjects. Figure 1 provides a summary of the analysis pipeline. Regularised regression with elastic net penalization (Zou & Hastie, 2005) was used to test the hypothesis that selected case-control differences in cortical morphometry were associated with variation in symptom severity (α =0.5, λ=100, k-fold cross-validation=10, n iterations =1000). In machine learning, we aim to use data to train a model to make accurate predictions on new, unseen or held-out datasets. Elastic net is an embedded technique that combines machine learning and feature reduction functions by implementing a regularization framework to obtain a reduced subset of selected features. Elastic net has been previously applied in machine learning for neuroimaging in Alzheimer’s disease classification and treatment outcome predictions in ADHD cohorts (Mwangi et al., 2014). In regularised regression, λ is a parameter controlling for the strength of regularization. The higher the value of λ, the more likely coefficients will be estimated towards zero with an increased penalty. α is the mixing parameter (0< α<1) and determines the relative quantities of L2 norm penalized regression (ridge regression) and L1 norm penalized regression (LASSO regularised penalization). Elastic net is an approach that combines both the L1 and L2 penalties of the LASSO and ridge methods.

A subset of the matched-pairs sample (33%) was held out as an out-of-sample test set independent of the subject data; that is these data were not used in the cross-validation steps (training set), and only examined as an independent validation of the model (test set). The remaining data was used as a training set to obtain optimal model weights for selected features. In the training set, we employed a strict k-fold cross-validation loop (10 folds, 1000 iterations) to train the model to predict differences in symptom severity between matched cases and controls in the out-of-sample test set. The model was trained within a k-fold cross-validation loop (k=10). The training set is first randomly partitioned into *k* subsamples of equal size. For each fold, one subsample is withheld for internal validation to test the model trained on *k*-1 subsamples. Each of the *k* subsamples were used as the validation set once per fold. Results from each fold were averaged to obtain a single estimation. Due to intrinsic randomness of model building, estimated coefficients may vary after each run. To account for stochastic error and to ensure robustness of estimates, the process was repeated for n=1000 iterations, and the averaged coefficient weights used to generate predictions in the out-of-sample test set.

Model goodness-of-fit was assessed by constructing a null distribution of symptom severity outcome. To generate a null distribution, the symptom severity (difference) variable was randomized across every sample observation of cortical thickness or surface area features using 5,000 permutations. For each iteration, model parameters were obtained using the same machine learning pipeline with regularised regression with elastic net penalty. The p-value of the initial model fit in the out-of-sample test set was computed as the proportion of iterations in the null distribution with model performance exceeding that of the initial model fit. Application and validation of recommended best practice for the regularised regression protocol are detailed elsewhere (Hendricks & Ahn, 2017; Vilares et al., 2017). The entire procedure was repeated for cortical thickness and surface area measures. Final model weights were obtained by fitting selected features on the entire dataset to allow independent model testing. To validate our approach against group-level prediction methods, we repeated the entire machine learning pipeline on the same matched ASD cohort but at the group-level without within-pair distance computations, adjusted for age, sex, site, IQ and intracranial volume effects.

Analyses were performed in the R environment (Team, 2014) using the *MatchIt* and *boot* wrapper tools (Ho et al., 2011; McArtor, Lubke, & Bergeman, 2016). Visualizations were generated with in-house scripts.

## Acknowledgements

Data analysis and interpretation was conducted within the Developmental Imaging research group, Murdoch Children’s Research Institute and the Children’s MRI Centre, Royal Children’s Hospital, Melbourne, Victoria. All procedures were conducted in accordance with the Declaration of Helsinki, and approved by The Royal Children’s Hospital Human Research Ethics Committee. The research was supported by the Murdoch Children’s Research Institute, The Royal Children’s Hospital, Department of Paediatrics The University of Melbourne and the Victorian Government’s Operational Infrastructure Support Program. The project was generously supported by RCH1000, a unique arm of The Royal Children’s Hospital Foundation devoted to raising funds for research at The Royal Children’s Hospital.

## Author contributions

EP designed the study, performed data analysis and visualizations and prepared the manuscript. GB contributed to the study design and manuscript preparation. CA contributed to data analysis and visualizations. SB and MS supervised the study and contributed to manuscript preparation.

## Footnotes

1 http://fcon_1000.projects.nitrc.org/indi/abide/

