## Supplementary materials for "Quantifying individual differences in brain morphometry underlying symptom severity in Autism Spectrum Disorders"

### Supplementary material

#### Supplementary Table 1. Mean beta coefficients of cortical thickness features selected by regularized regression with elastic net penalty

| Cortical thickness features | Hemisphere | Mean | SD | Lower bound | Upper bound |
| --- | --- | --- | --- | --- | --- |
| Inferior temporal gyrus | R | 0.49 | 0.022 | 0.45 | 0.54 |
| Pars orbitalis | L | 0.26 | 0.0058 | 0.25 | 0.27 |
| Cuneus cortex | R | 0.23 | 0.0059 | 0.22 | 0.24 |
| Inferior parietal lobule | R | 0.19 | 0.012 | 0.16 | 0.21 |
| Insula | R | 0.15 | 0.0090 | 0.13 | 0.17 |
| Isthmus cingulate gyrus | R | 0.15 | 0.0066 | 0.14 | 0.17 |
| Medial orbitofrontal | R | 0.11 | 0.0050 | 0.10 | 0.12 |
| Posterior cingulate gyrus | R | 0.10 | 0.0021 | 0.099 | 0.11 |
| Caudal anterior cingulate gyrus | R | 0.098 | 0.0052 | 0.089 | 0.11 |
| Rostral middle frontal gyrus | L | 0.00069 | 0.0022 | 0.00 | 0.0093 |
| Pericalcerine cortex | R | -0.049 | 0.0037 | -0.057 | -0.042 |
| Pars triangularis | R | -0.052 | 0.0015 | -0.055 | -0.049 |
| Fusiform gyrus | L | -0.062 | 0.0029 | -0.068 | -0.056 |
| Middle temporal gyrus | L | -0.082 | 0.0014 | -0.085 | -0.080 |
| Lingual gyrus | L | -0.14 | 0.0066 | -0.15 | -0.13 |
| Entorhinal cortex | R | -0.15 | 0.0083 | -0.17 | -0.13 |
| Inferior temporal gyrus | L | -0.21 | 0.014 | -0.24 | -0.19 |
| Fusiform gyrus | R | -0.24 | 0.0039 | -0.25 | -0.23 |
| Middle temporal gyrus | R | -0.36 | 0.018 | -0.40 | -0.33 |

#### Supplementary Table 2. Mean beta coefficients of surface area features selected by regularized regression with elastic net penalty

| Surface area features | Hemisphere | Mean | SD | Lower bound | Upper bound |
| --- | --- | --- | --- | --- | --- |
| Isthmus cingulate gyrus | L | 0.23 | 0.0079 | 0.21 | 0.24 |
| Caudal middle frontal gyrus | L | 0.15 | 0.0063 | 0.13 | 0.15 |
| Supramarginal gyrus | L | 0.13 | 0.0038 | 0.12 | 0.13 |
| Entorhinal cortex | R | 0.11 | 0.0028 | 0.11 | 0.16 |
| Isthmus cingulate gyrus | R | -0.14 | 0.0055 | -0.15 | -0.13 |
| Rostral anterior cingulate gyrus | R | -0.15 | 0.00012 | -0.15 | -0.15 |
| Rostral anterior cingulate gyrus | L | -0.15 | 0.0029 | -0.15 | -0.14 |
| Inferior parietal lobule | L | -0.15 | 0.0057 | -0.16 | -0.14 |
| Rostral middle frontal gyrus | L | -0.21 | 0.0069 | -0.21 | -0.19 |
| Insula | L | -0.24 | 0.0052 | -0.24 | -0.23 |

#### Supplementary Table 3. Comparison of model fit in subject-level, distance-based model to the group-level model

| **Model** | **R^2^** | **Mean Squared Error** | **Correlation, *r*** |
| --- | --- | --- | --- |
| **Cortical Thickness** |  |  |  |
| Subject-level model | 0.15 | 0.98 | 0.39 |
| Group-level model | 0.0000434 | 1.025 | -0.00659 |
| **Surface Area** |  |  |  |
| Subject-level model | 0.18 | 1.01 | 0.42 |
| Group-level model | 0.0158 | 1.009 | 0.13 |

*Note. Cortical features selected by our approach (subject-level, distance-based model) significantly outperformed common methods (group-level modelling) in predicting symptom severity in independent out-of-sample validation testing. Both models were trained and evaluated using the same rigorous machine learning pipeline.*

#### Supplementary Figure 1. Out-of-sample predictions for cortical thickness and surface area features without subject-level modelling.

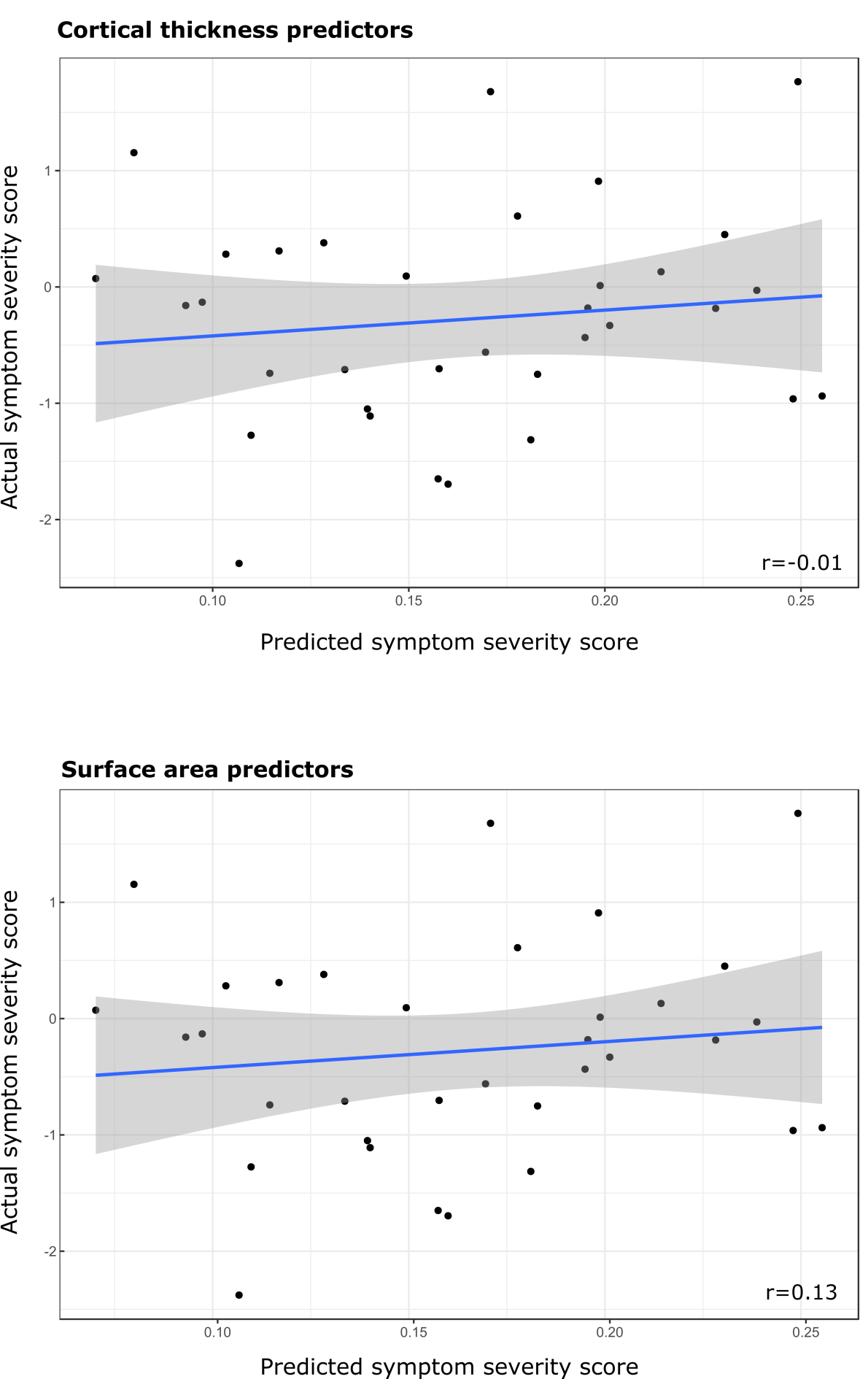

***Supplementary Figure 1. Out-of-sample predictions for cortical thickness and surface area features without subject-level modelling.*** *Regularized regression models for cortical thickness and surface area features trained on data without subject-level information performed poorly in predicting actual symptom severity scores in independent out-of-sample validation.*
